# Directed yeast genome evolution by controlled introduction of trans-chromosomic structural variations

**DOI:** 10.1101/2021.07.26.453910

**Authors:** Bin Jia, Jin Jin, Ming-Zhe Han, Bing-Zhi Li, Ying-Jin Yuan

## Abstract

Naturally occurring structural variations (SVs) are a considerable source of genomic variation and can reshape chromosomes 3D architecture. The synthetic chromosome rearrangement and modification by loxP-mediated evolution (SCRaMbLE) system has been proved to generate random SVs to impact phenotypes and thus constitutes powerful drivers of directed genome evolution. However, how to reveal the molecular mechanism insights into the interactions between phenotypes and complex SVs, especially inversions and translocations, has so far remained challenging. In this study, we develop a SV-prone yeast strain by using SCRaMbLE with two synthetic chromosomes, synV and synX. An heterologous biosynthesis pathway allowing a high throughput screen for increased yield of astaxanthin is used as readout and a proof of concept for the application of SV in industry. We report here that complex SVs, including a pericentric inversion and a trans-chromosomes translocation between synV and synX, result in two neochromosomes and a 2.7-fold yield of astaxanthin. We demonstrated that inversion and inversion reshaped chromosomes 3D architecture and led to large reorganization of the genetic information nearby the breakpoint of the SVs along the chromosomes. Specifically, the pericentric inversion increased the expression of STE18 and the trans-chromosomic translocation increased the expression of RPS5 and MCM22, which contributed to higher astaxanthin yield. We also used the model learned from the aforementioned random screen and successfully harnessed the precise introduction of trans-chromosomes translocation and pericentric inversions by rational design. Overall, our work provides an effective tool to not only accelerate the directed genome evolution but also reveal mechanistic insight of complex SVs for altering phenotypes.

## Introduction

Chromosomes are highly dynamic objects that often undergo large structural variations(SVs), which in turn can reshape chromosomes and impact phenotypes(Pevzner and Tesler 2003; Dujon et al. 2004; Redon et al. 2006; Darling et al. 2008; Kidd et al. 2008; Conrad et al. 2010; Dujon 2010; Pang et al. 2010; Henssen et al. 2017; Yue et al. 2017; Peter et al. 2018). Naturally occurring SVs comprised both unbalanced SVs that are copy number variations including deletions and duplications, and balanced SVs that are copy number neutral including inversions and translocations(Fleiss et al. 2019). Both the unbalanced SVs and balanced SVs can modify the 3D architecture of the chromosomes and potentially affect genetic functioning, which contributed to the genome evolution. For example, Selmecki.et al showed tetraploids yeast undergo significantly faster adaptation in a poor carbon source environment with larger deletion of chromosomes. (Selmecki et al. 2015). Li.et al found that deletion of left arm of synthetic chromosome X and duplication of chromosome VIII (trisomy) lead to increased rapamycin resistance in yeast(Li et al. 2019). With the development of genome medicine, it is indicated reciprocal translocation between chromosomes have contributed to several human diseases, for instance, chronic myeloid leukemia (chromosomes 9 with 22) and Mobius-like syndrome (chromosomes 1 with 2)(Eyupoglu et al. 2016) (Nishikawa et al. 1997). In addition, pericentric inversion of chromosome 9 is one of the most common chromosomal abnormalities and can be associated with ambiguous genitalia in children(Sotoudeh et al. 2017). It is challenge to map the balanced SVs than unbalanced SVs due to the copy number neutral. So far, the detailed molecular mechanism underlying balanced SVs remained unexplored due to lack of cell models.

The baker yeast (Saccharomyces cere*visiae*) appreciated for the availability of its powerful genetic tools, biosafety and fast growth rate has been used extensively in synthetic biology for industrial purposes. High-throughput phenotypic screenings have been employed to identify genome-wide deletions and duplications that increase the yield of yeast synthetic biology products. The completion of the yeast gene-knockout collection (YKOC) has enabled the analysis of synthetic genetic interactions(Puddu et al. 2019). The construction of segmental duplications covering the whole *S. cerevisiae* genome has been used to produce strains with enhanced tolerance to stresses(Natesuntorn et al. 2015). Many rational design strategies, including overexpression and downregulation, are time-consuming because identifying gene targets remains an challenge due to the complexity of genetic networks (Si et al. 2015)[Rapid prototyping of microbial cell factories via genome-scale engineering. Biotechnol Adv 2015, 33:1420-1432.]. Many successful examples demonstrate that the synthetic yeast chromosome could be reprogrammed for various purposes with greatly enhanced metabolic capacity(Jia et al. 2018; Shen et al. 2018; Wu et al. 2018). The Synthetic Chromosome Rearrangement and Modification by LoxP-mediated Evolution (SCRaMbLE) technology have been developed to generate larger-scale SVs(Blount et al. 2018; Jia et al. 2018; Liu et al. 2018; Luo et al. 2018; Shen et al. 2018; Wang et al. 2018; Wu et al. 2018; Li et al. 2019; Ma et al. 2019; Gowers et al. 2020). SCRaMbLE is a powerful approach to study how chromosomal architecture impacts phenotypes and can be to mimic the Darwinian evolution process in the test tube for targeted phenotypes. Thus, SCRaMbLE can be viewed as natural molecular tools for directed genome evolution by iterative rounds of mutagenesis and screening or selection on the genome scale. So far, SCaMbLE has been used for directed genome evolution mainly with signal synthetic chromosome.

In this work we report that one by using SCaMbLE and two synthetic chromosomes, can for the first time creates chromosome inversion and translocation events on a larger scale and at a higher frequency. Introducing the same yeast cell with two synthetic chromosomes, each containing hundreds of non-directional loxPsym sites positioned downstream of non-essential genes and major landmarks provides a novel approach to genome reorganization especially SVs. We evaluate the effects of SVs on the enhanced production of an exogenous but commercially valuable terpenoid, astaxanthin. Two cycles of SCRaMbLE were used to iteratively generate complex SVs and improve the astaxanthin yield. We used the high-resolution chromosome conformational capture (Hi-C) sequencing approach to investigate the 3D conformations of chromosomes with complex SVs(Akdemir et al. 2020), and observed an inversion spanning the centromere and a translocation between synV and synX. We developed a method to precisely reconstitute the translocations and inversions, and recaptured their combined effects on increasing the yield of astaxanthin by 2.7-fold. Our work thus provides a new toolkit for directed evolution of genomic assembly and reorganization.

## Results

### SCRaMbLE with two synthetic chromosomes for directed genome evolution

To investigate more complex SVs and biosynthesis pathways, we used the haploid strain SYN510 containing both synV with 176 loxPsym sites(Xie et al. 2017) and synX with 245 loxPsym sites(Wu et al. 2017c) for the biosynthesis of astaxanthin. Astaxanthin is the end metabolite of the MVA pathway; it has tremendous antioxidant activity and is widely used in nutraceuticals, cosmetics and aquaculture(Ambati et al. 2014; Igielska-Kalwat et al. 2015). The astaxanthin biosynthesis pathway consists of 5 genes. *CrtE*, *CrtI* and *CrtYB* are involved in carotenoid biosynthesis, after which a reticular metabolic pathway with β-carotene ketolase (CrtW) and β-carotene hydroxylase (CrtZ) performs a two-step hydroxylation and ketolation reaction(Supplementary information, Fig. 1a). Our previous study demonstrated that the combination of Aa_CrtZ–BDC263_CrtW showed lower intermediate accumulation and achieved better astaxanthin yield(Wang et al. 2017). To increase the stability of the astaxanthin biosynthesis pathway, all five genes were assembled together and integrated at the *YEL063C*/*CAN1* locus of synV to generate the yeast strain YJJ001 (Fig. 1a). It has been reported that the pCRE4 plasmid (pGal1-Cre-EBD-tCYC1) has tight control of Cre recombinase and can be used for Multiplex SCRaMbLE Iterative Cycling (MuSIC). Thus, the pCRE4 plasmid with a Ura3 marker was transformed into YJJ001 for the control of SCRaMbLE.

**Fig. 1.**
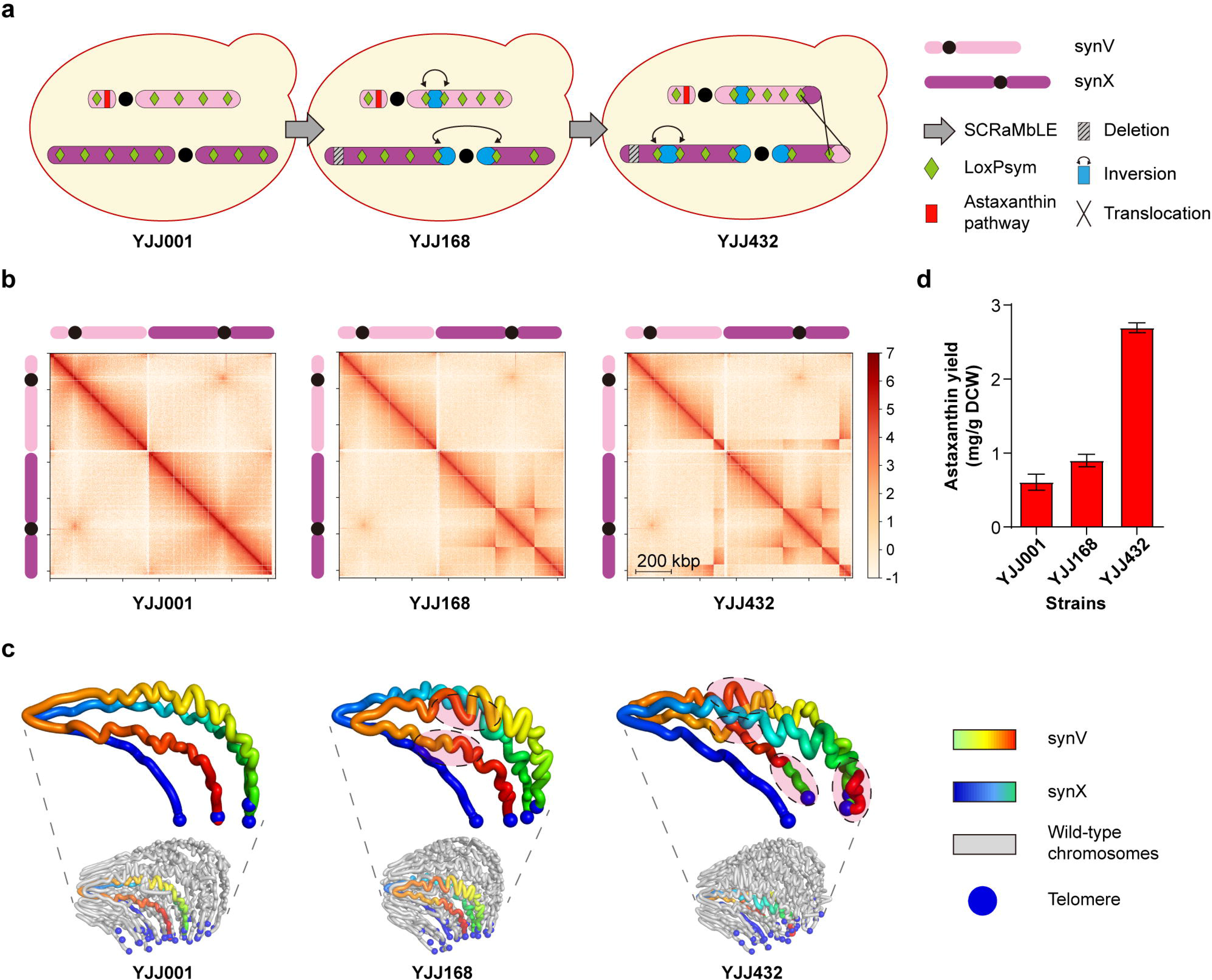
SCRaMbLE with two synthetic chromosomes generating complex SVs. a, Schematic representation of structural variations in YJJ001, YJJ168 and YJJ432. Synthetic chromosome V and synthetic chromosome X are abbreviated as synV and synX, respectively. b, Normalized contact maps (bin size, 2 kb) of synthetic chromosomes in three different strains: YJJ117, YJJ168 and YJJ432. All Hi-C reads are mapped against the reference genome of the parental strain YJJ001. c, Three-dimensional model of the YJJ001, YJJ168 and YJJ432 genomes. Synthetic chromosomes and wild chromosomes are indicated with graduated color and gray, respectively. The SV area are indicated with the dashed oval. d, The astaxanthin yield of control strain YJJ001 and SCRaMbLEd strains YJJ168 and YJJ432.

In the first cycle of SCRaMbLE, strain YJJ168 was selected from the SCRaMbLEd yeast pool due to its dark red pigmentation (Supplementary information, Fig. 1b). To generate more complicated SVs and the desired phenotype, we used YJJ168 for the second cycle of SCRaMbLE to yield another SCRaMbLEd strain, YJJ432. To identify the chromosome SVs in haploids caused by SCRaMbLE, we deep sequenced YJJ168 and YJJ432. As shown in Fig. 1a and Supplementary information, Fig. 2, a 3 kb deletion (YJL218W-YJL217W), a 1 kb inversion (YER044C) and a 210 kb pericentric inversion (YJL052C-A-YJR071C) with 127 genes were observed in the SCRaMbLEd strain YJJ168. It is interesting that the upstream and downstream of inversions of YJL052C- A-YJR071C were on the left arm and right arm of synX, respectively. In other words, this inversion spanned the centromere of synX. As an iterated SCRaMbLEd strain from YJJ168, YJJ432 was observed to contain not only all the same SVs as YJJ168 but also a inversion of YJL170C-YJL158C and a trans-chromosomictranslocation. It is noted that the trans-chromosomictranslocation caused an exchange of 59 kb from the right arm of synV and 74 kb from the right arm of synX that had not previously been reported. From the sequencing data above, we find that the second-generation strains can inherit genotypic SVs from first-cycle SCRaMbLEd strains. PCR verification of novel junctions also confirmed that inversion and translocation events occurred in YJJ168 and YJJ432 (Supplementary information, Fig. 3).

**Fig. 2.**
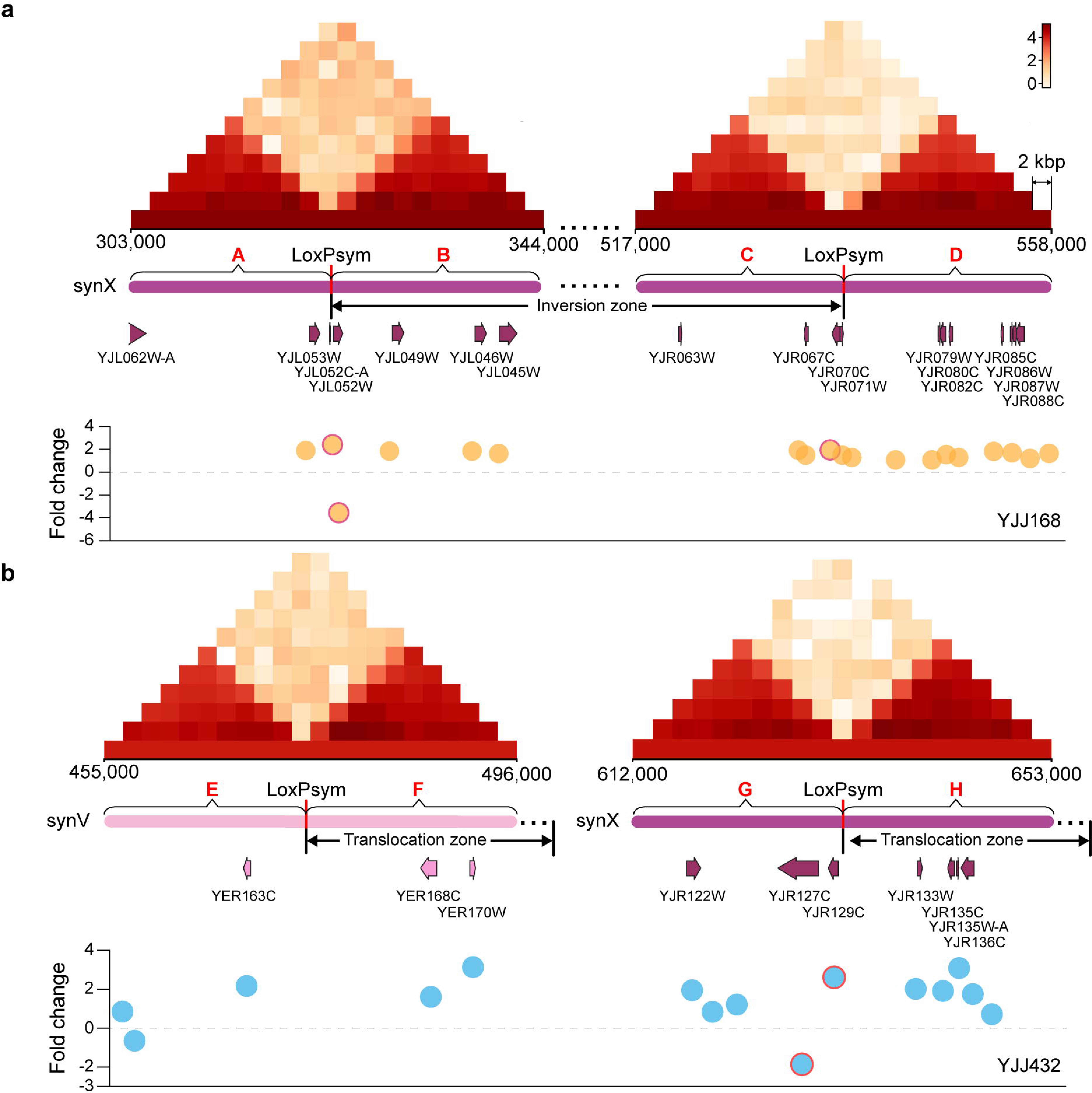
Quantified interactions of loxP breakpoint-flanking regions. a, Triangle heat maps representing the contact frequencies among 20 bin regions (length 20 kb) upstream and downstream of two recombination sites (YJL052C-A and YJR071C) in YJJ168, labeled A, B, C, and D. Dots show fold changes in expression in the A, B, C, and D regions. b, Triangle heat maps representing the contact frequencies among 20 bin regions (length 20 kb) upstream and downstream of the two recombination sites (YER164W and YJR130C) in YJJ432, labeled E, F, G, and H. Dots show fold changes in expression in the E, F, G, H regions.

**Fig. 3.**
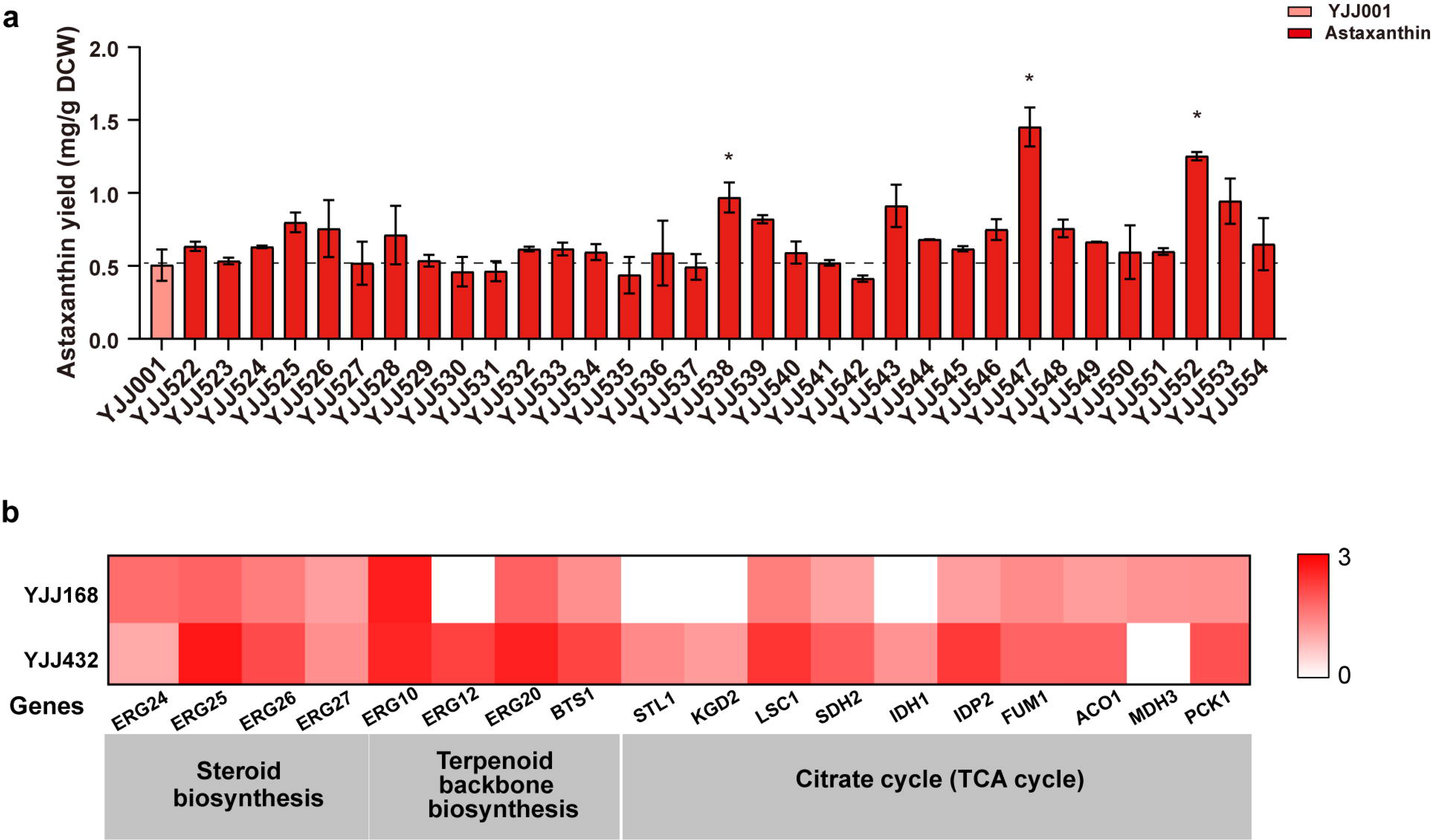
a, The astaxanthin yield of control strain YJJ001 and genes overexpression and deletion around pericentric inversion strain breakpoint-flanking regions and genes overexpression and deletion around trans-chromosomic translocation strain breakpoint-flanking regions. (Student’s t-test; *P < 0.05). b, Enrichment analysis of transcriptomics of YJJ168 and YJJ432 were used to identify differentially transcribed genes with known functions involved in KEGG pathways.

To investigate the structural organization of the chromosome SVs, Hi-C contact maps (Fig. 1b) and 3D representations (Fig. 1c) of synV and synX were generated from YJJ001, YJJ168 and YJJ432. The contact maps of synV and synX in YJJ168 and YJJ432 showed several new interactions, indicating large-scale rearrangements in both SCRaMbLEd strains. YJJ168 exhibited a new interaction indicating a large inversion (YJL052C-A-YJR071C) event. The contact map of YJJ432 revealed three new interactions caused by the large inversion (YJL052C-A-YJR071C), one inversion (YJL170C-YJL158C) in synX and one translocation between the right arm of synV (YJR130C) and the right arm of synX (YER164W). A three-dimensional model of the 16 yeast chromosomes and amplified chromosomes of synV and synX are provided in Fig. 1c. The centromeres gather at the spindle pole, and all chromosomes are clustered by the centromeres at one pole of the nucleus. The telomeres are clustered at the other end. The 3D representations show that the average trajectories of chromosomes in YJJ168 and YJJ432 did not appear to be substantively altered compared with those in YJJ001, with the synthetic chromosomes neighboring the same chromosomes as the native ones. New connections were generated in synX of strain YJJ168 and strain YJJ432 due to the inversion spanning the centromere. In strain YJJ432, the right arms of synV and synX generate two new connections because of the translocation between synV and synX (the area within the dashed oval). It is indicated that the translocation on the right arm of synV has a closer spatial distance with synX and consequently has stronger interactions.

Meanwhile, the two SCRaMbLEd strains and YJJ001 were subjected to fermentation experiments in YPD medium, and the astaxanthin yield was analyzed by high-performance liquid chromatography (HPLC). As illustrated in Fig. 1d, the astaxanthin yield in strain YJJ001 was 0.61 mg/g DCW, and the astaxanthin yield of the two SCRaMbLEd strains (YJJ168 and YJJ432) was increased, to 0.90 and 2.7 mg/g DCW, respectively. The astaxanthin yield was increased 1.48- to 4.43-fold in the SCRaMbLEd strains compared with the nonSCRaMbLEd parent strain. These results revealed the new interactions of synthetic chromosomes in SCRaMbLEd strains via Hi-C technology. SCRaMbLE strains provide sufficiently detectable SVs, indicating that the SCRaMbLE system provides a driving force to alter cell phenotypes through rearranging gene order and number.

### Reorganization of the genetic information on the regions of SVs

The pericentric inversion and trans-chromosomic translocation alter the spatial positions of the breakpoint-flanking regions, which may which may led to large reorganization of the genetic information along the chromosomes. To explore whether the genetic information in the breakpoint-flanking regions were affected, we examined transcription profile on the 20 kb regions upstream and downstream of the two loxPsym breakpoints. As shown in Fig. 2a, the upstream and downstream regions of the breakpoints of the pericentric inversion in YJJ168 were named A, B, C, and D, respectively. We first observed changes in the transcription levels of YJL053W, YJL052C-A and YJL052W near the left breakpoint and the transcription levels of YJL067C, YJL070C and YJL071W near the right breakpoint. In the whole 80 kb region consisting of A, B, C and D, 19 of 51 genes showed changes in expression compared with the control strain YJJ001(Log2FoldChange > 1-fold), with 1 gene of YJL052W downregulated and 18 genes (YJL053W, YJL052C-A, YJL049W, YJL046W, YJL045W, YJR067C, YJR068W, YJR070C, YJR071W, YJR072C, YJR076C, YJR079W, YJR080C, YJR082C, YJR085C, YJR086C, YJR087W and YJR088C) upregulated, respectively. Around the left breakpoint, the changes in YJL052C-A and YJL052W were the most obvious, whereas around the right arm recombination site, the change in YJR070C was the most obvious. Similarly, the regions upstream and downstream of the breakpoint in the right arm of synV and the breakpoint in the right arm of synX in YJJ432 were labeled E, F, G, and H, respectively (Fig. 2b). Compared with the control strain, 14 of the 38 genes in these regions exhibited changed expression(Log2FoldChange > 1-fold), with 2 genes (YER158C and YJR127C) downregulated and 12 genes (YER156C, YER163C, YER168C, YER170W, YJR122W, YJR123W, YJR125C, YJR129C, YJR133W, YJR135C, YJR135W-A, YJR136C) upregulated, respectively. The results indicated that the expression of genes in the breakpoint-flanking regions was indeed affected by different types of SVs. To further quantify the interactions of the recombination sites, we counted the interaction values of the four points on the interaction matrix of YJJ168 and YJJ432 and counted the differences between different interaction domains (areas marked by boxes in Supplementary information, Fig. 4). As the results show, regions A and C and regions B and D had significant interactions in inversion strains YJJ168 and YJJ432. In translocation strain YJJ432, the interactions of regions E and H and F and G were increased as a result of the greater proximity of the right arm of synV (YJR130C) to synX and the right arm of synX (YER164W) to synV. These results further indicate that two sites that are closer in space have greater interaction. It is spectacular that the pericentric inversions and trans-chromosomes translocation changed larger number of genes position and the transcription direction, which affected the transcription levels of genes in the regions nearby the breakpoint of SVs(Wu et al. 2017b).

**Fig. 4.**
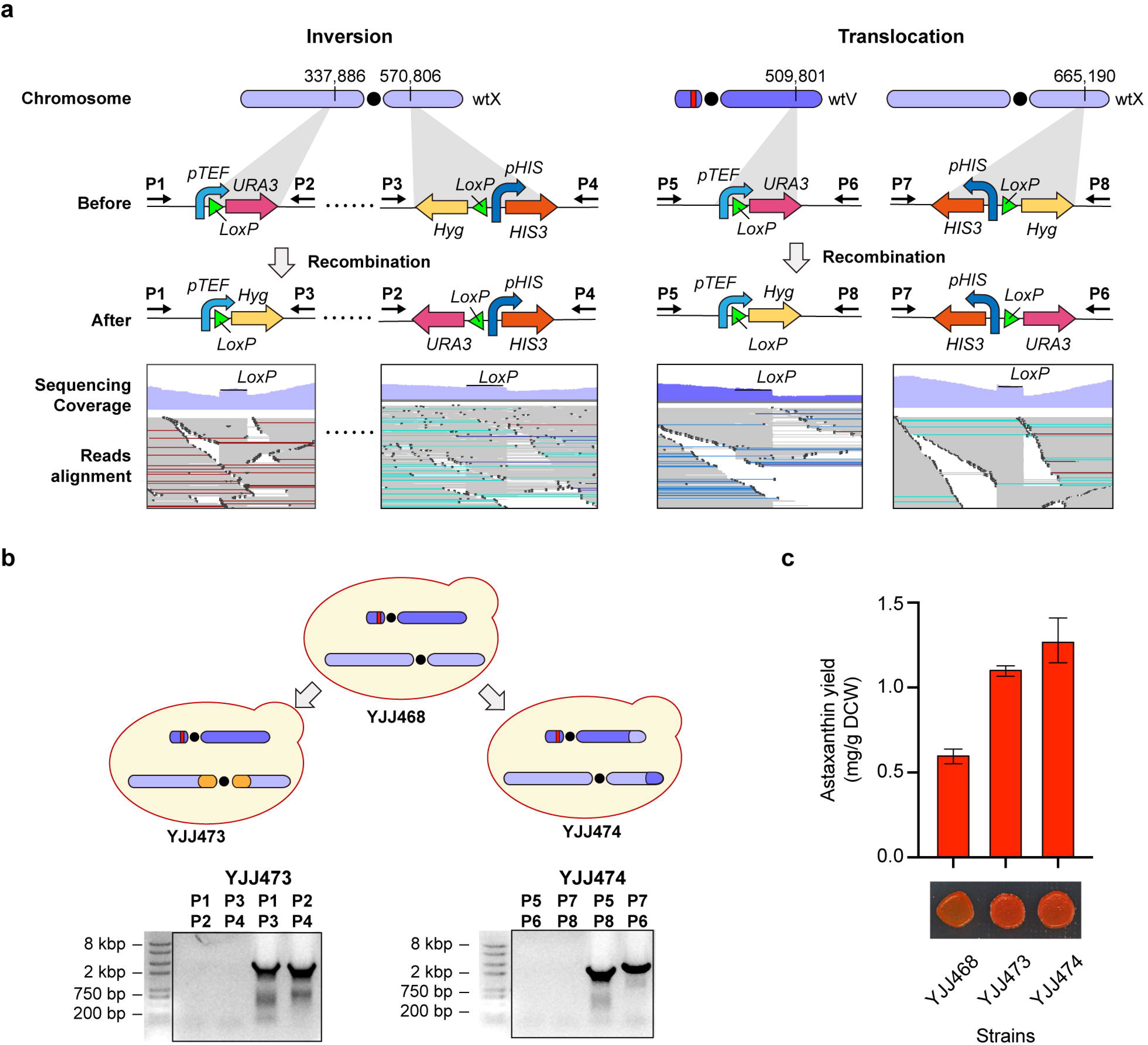
Rational translocation and inversion of chromosomes. a, The process of inversion and translocation was validated in wild-type strains. Wild-type chromosome V and wild-type chromosome X are abbreviated as wt V and wt X, respectively. Validation strains exhibited inversion and translocation events between two LoxP sites. These events were identified by aligning all raw reads. b, Verification of inversion and translocation by junction PCR. c, The astaxanthin yield of the wild-type control strain YJJ468, inversion strain YJJ473 and translocation strain YJJ474. Serial dilution assay of YJJ468, YJJ473, YJJ474.

### Mapping of genetic targets contributing to higher astaxanthin yield

To identify the genetic targets with regard to the improvement of astaxanthin yield, the above down-regulated genes and up-regulated genes in the regions nearby the breakpoint of SVs were knocked out and overexpressed in the strain YJJ001 for fermentation assay, respectively(Table 1). As illustrated in Fig. 3a, compared to the control strain, astaxanthin yields were significantly increased in strains with YJR086W, YJR123W and YJR135C over-expression strain. The function of these three genes are briefly described in Table 2. Although the exact reason for these genes to be beneficial to astaxanthin yield remains to be elucidated, it is possible to speculate the mechanism resulting in the favorable changes of this phenotype from functions previously described in the literature. The result indicated upregulated these genes have contributed to higher astaxanthin yield.

**Table 1.** *S. cerevisiae* strains used in this study

| Strains | Description | Sources |
| --- | --- | --- |
| BY4741 | *MATa, HIS3Δ1, LEU2Δ0, MET15Δ0, URA3Δ0* | (Brachmann et al. 1998) |
| SYNVX | *MATa, HIS3Δ1, LEU2Δ0, MET15Δ0, URA3Δ0* | This study |
| YJJ001 | yXZX573, CAN1:: astaxanthin pathway with Leu2 marker | This study |
| YJJ002 | Introducing plasmid pGAL1-Cre-EBD-GFP-tCYC1 into strain YJJ001 | This study |
| YJJ168 | SCRaMbLEd strain from the YJJ002 | This study |
| YJJ432 | SCRaMbLEd strain from the YJJ168 | This study |
| YJJ468 | Astaxanthin producing strain, BY4741 | This study |
| YJJ473 | YJJ468 with inversion (YJL052C-A-YJR071C) strain | This study |
| YJJ474 | YJJ468 with translocation (YJR130C and YER164W) strain | This study |
| YJJ522 | YJJ001 with YJL053W overexpression | This study |
| YJJ523 | YJJ001 with YJL052C-A overexpression | This study |
| YJJ524 | YJJ001 with YJL052W deletion | This study |
| YJJ525 | YJJ001 with YJL049W overexpression | This study |
| YJJ526 | YJJ001 with YJL046W overexpression | This study |
| YJJ527 | YJJ001 with YJL045W overexpression | This study |
| YJJ528 | YJJ001 with YJR067C overexpression | This study |
| YJJ529 | YJJ001 with YJR068W overexpression | This study |
| YJJ530 | YJJ001 with YJR070C overexpression | This study |
| YJJ531 | YJJ001 with YJR071W overexpression | This study |
| YJJ532 | YJJ001 with YJR072C overexpression | This study |
| YJJ533 | YJJ001 with YJR076C overexpression | This study |
| YJJ534 | YJJ001 with YJR079W overexpression | This study |
| YJJ535 | YJJ001 with YJR080C overexpression | This study |
| YJJ536 | YJJ001 with YJR082C overexpression | This study |
| YJJ537 | YJJ001 with YJR085C overexpression | This study |
| YJJ538 | YJJ001 with YJR086W overexpression | This study |
| YJJ539 | YJJ001 with YJR087W overexpression | This study |
| YJJ540 | YJJ001 with YJR088C overexpression | This study |
| YJJ541 | YJJ001 with YER156C overexpression | This study |
| YJJ542 | YJJ001 with YER158C deletion | This study |
| YJJ543 | YJJ001 with YER163C overexpression | This study |
| YJJ544 | YJJ001 with YER168C overexpression | This study |
| YJJ545 | YJJ001 with YER170W overexpression | This study |
| YJJ546 | YJJ001 with YJR122W overexpression | This study |
| YJJ547 | YJJ001 with YJR123W overexpression | This study |
| YJJ548 | YJJ001 with YJR127 deletion | This study |
| YJJ549 | YJJ001 with YJR125C overexpression | This study |
| YJJ551 | YJJ001 with YJR133W overexpression | This study |
| YJJ552 | YJJ001 with YJR135C overexpression | This study |
| YJJ553 | YJJ001 with YJR135W-A overexpression | This study |
| YJJ554 | YJJ001 with YJR136C overexpression | This study |

**Table 2.**
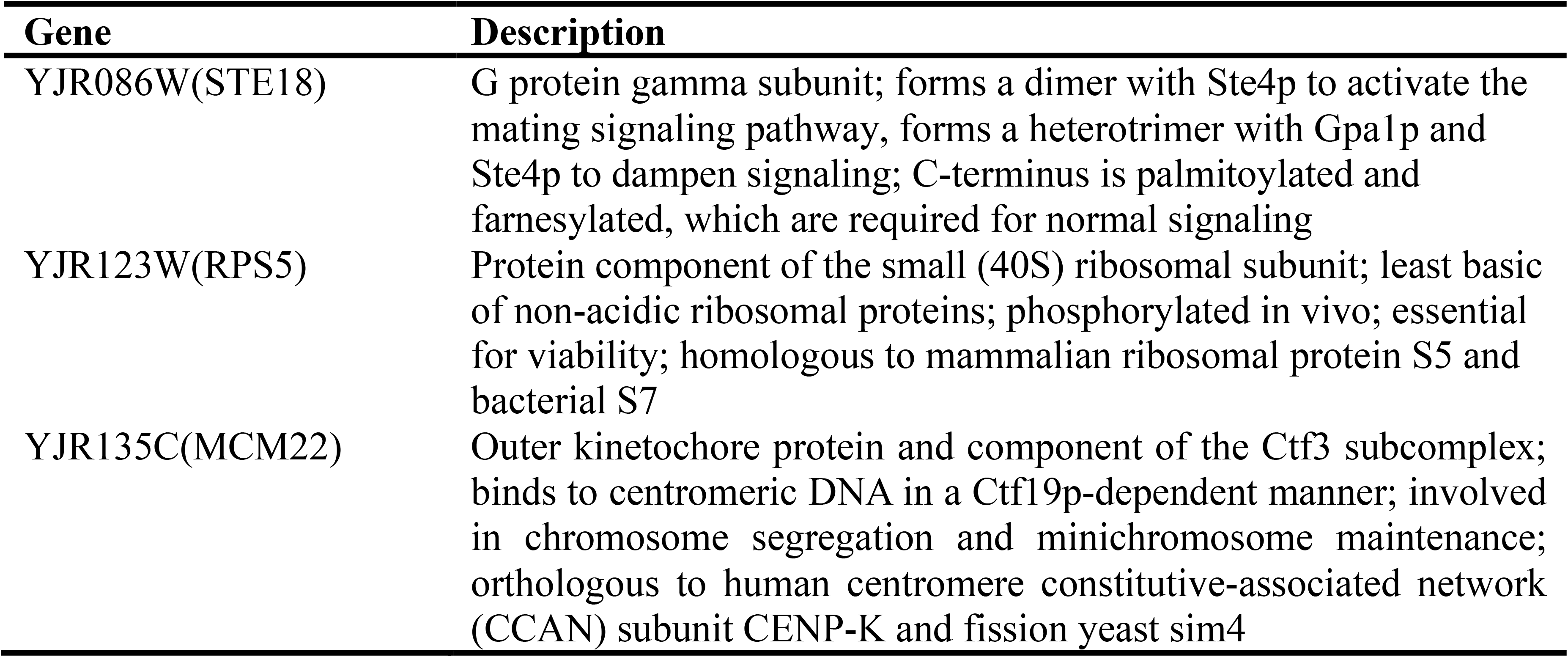
Known gene functions for beneficial over-expression

| Gene | Description |
| --- | --- |
| YJR086W(STE18) | G protein gamma subunit; forms a dimer with Ste4p to activate the mating signaling pathway, forms a heterotrimer with Gpa1p and Ste4p to dampen signaling; C-terminus is palmitoylated and farnesylated, which are required for normal signaling |
| YJR123W(RPS5) | Protein component of the small (40S) ribosomal subunit; least basic of non-acidic ribosomal proteins; phosphorylated in vivo; essential for viability; homologous to mammalian ribosomal protein S5 and bacterial S7 |
| YJR135C(MCM22) | Outer kinetochore protein and component of the Ctf3 subcomplex; binds to centromeric DNA in a Ctf19p-dependent manner; involved in chromosome segregation and minichromosome maintenance; orthologous to human centromere constitutive-associated network (CCAN) subunit CENP-K and fission yeast sim4 |

To further explore the global affection of chromosomal rearrangements on metabolic pathways, enrichment analysis of transcriptomics was used to identify differentially transcribed genes with known functions involved in KEGG pathways. Several genes involved in terpenoid backbone biosynthesis, ergosterol biosynthesis and citrate cycle were upregulated in the YJJ168 and YJJ432 (Fig. 3b). In SCRaMbLEd strain YJJ168, the upregulated genes ERG24, ERG25, ERG26 and ERG27 were involved in ergosterol biosynthesis, while ERG10, ERG20 and BTS1 were involved in terpenoid backbone biosynthesis. In YJJ432, ERG24, ERG26 and ERG27, which are involved in the biosynthesis of ergosterol, and ERG10, ERG12, ERG20 and BTS1, which are involved in terpenoid backbone biosynthesis, were found to be upregulated. As a consequence, these two pathways could increase the flux of the MVA pathway and cell storage capacity of hydrophobic carotenoids(Gao et al. 2017; Wu et al. 2017a). The improvement of citrate cycle could provide sufficient reducing power and energy for the MVA pathway. These transcriptome analyses indicated that the improved yield of astaxanthin was caused by the upregulation of these genes, further clarifying the direct effect of the SVs on astaxanthin accumulation. The results demonstrated the SVs can led to large reorganization of the genetic information along the chromosomes and accelerate the directed genome evolution with selection of targeted phenotypes.

### Rational introduction of inversion and translocation in yeast

To verify the effect of the pericentric inversion and trans-chromosomic translocation, we developed a method to rationally generate these inversions and translocations. First, we constructed an astaxanthin-producing strain named YJJ468 from the wild-type strain BY4741 by integrating the biosynthesis pathway of astaxanthin at the *YEL063C*/*CAN1* locus of chromosome V. Then, we designed an on/off switch consisting of two markers (URA3 and Hyg). As shown in Fig. 4a, the URA3 promoter was inserted upstream of loxP site 1, and the open reading frame of URA3 was inserted downstream of loxP site 1, allowing the expression of URA3. Hyg without a promoter was positioned immediately downstream of loxP site 2. A complete HIS3 marker was positioned upstream of loxP site 2 as a selection marker to help loxP-Hyg integrate into the chromosome. The pGal1-Cre-EBD-tCYC1 plasmid with a G418 marker was transformed into strain YJJ468 (Supplementary Fig. 7). The activation of Cre recombinase will catalyze the recombination of the two loxP sites, leading to a change in the phenotype of the yeast from Hyg- to Hyg+. Consequently, hygromycin media could be used to select translocation and inversion strains. Using the above approach, strain YJJ473 with pericentric inversion and strain YJJ474 with trans-chromosomic translocation were generated and isolated by screening on SC-hygromycin agar plates. Strains YJJ473 and YJJ474 were further analyzed using whole-genome sequencing to verify the genomic variations produced by the recombination. As Fig. 4a shows, the inversion and translocation region had breakpoints at the loxP sites in YJJ473 and YJJ474, indicating that the desired inversion and translocation were artificially recreated from the wild-type strain YJJ468.

Meanwhile, PCR was used to further verify the inversion and translocation. Before recombination, sequences in YJJ468 could be amplified by primer pairs 1/2, 3/4, 5/6, and 7/8. After recombination, sequences in YJJ473 could be amplified by primer pairs 1/3 and 2/4, while sequences in YJJ474 could be amplified by primer pairs 5/8 and 6/7 (Fig. 4b). Therefore, we can confirm that inversion and translocation events occurred in YJJ473 and YJJ474. Compared to the initial strain yJJ468, YJJ473 and YJJ474 had enhanced astaxanthin production phenotypes, yielding 1.11 mg/g DCW and 1.28 mg/g DCW, respectively, which were similar to those of the SCRaMbLEd strains (Fig. 4c). YJJ473 and YJJ474 increased the astaxanthin yield 1.8- and 2.1-fold relative to the parent strain YJJ468. These results indicated that the YJL052C-A-YJR071C inversion and YJR130C-YER164W translocation increased the production of astaxanthin and were responsible for the enhanced astaxanthin production phenotypes of SCRaMbLE strains yJJ168 and yJJ432. The results indicated that the YJL052C-A-YJR071C inversion and YJR130C-YER164W translocation contributed to the improvement of astaxanthin production. It is anticipated that we can used the model learned from the random screen of SCRaMbLE and successfully harnessed the precise introduction of pericentric inversions and trans-chromosomic translocation by rational design.

## Discussion

Our study focused on investigating complex chromosome SVs by enabling haploid yeast containing multiple synthetic chromosomes to be SCRaMbLEd, which has significant advantages over pre-existing methods for laboratory evolution. First, traditional evolution approaches, such as physical mutagenesis or chemical mutagenesis, typically cause hundreds of single nucleotide polymorphisms (SNPs) or insertions and deletions (indels)(Jin et al. 2018), which make it difficult to confirm the targets responsible for observed changes in phenotypes. In comparison, SCRaMbLE has been demonstrated to rapidly generate larger scale SVs, providing clear targets to verify and understand more deeply. In this study, we have demonstrated that pathway- doubling is not the only way to increase yield (Jia et al. 2018), changes in map position and direction of endogenous genes can also affect yield. Second, the synthetic yeast project (Sc2.0) aims to synthesize 16 yeast chromosomes, each of which contains hundreds of loxPsym sites, downstream of every nonessential gene. Our research has shown that SCRaMbLE with two synthetic chromosomes can generate translocations between two synthetic chromosomes. Thus, it is anticipated that the integration of multiple synthetic chromosomes will extend the diversity and quantity of SVs. Moreover, SCRaMbLE with multiple synthetic chromosomes provided a powerful model in which to investigate the trajectory of chromosome SVs during iterative evolution. In this study, YJJ432 not only inherited the inversion spanning the centromere from YJJ168 but also acquired new genetic changes, namely, a translocation between two synthetic chromosomes.

The SCRaMbLE system provided sufficiently large structural variations to allow us to analyze novel interactions and structures via Hi-C technology. The translocation strain YJJ474 on the right arm of synV (YJR130C) has a closer spatial distance with synX and consequently has stronger interactions. Chromosomal SVs cause changes in the 3D architecture of the genome and potentially alter cellular functions. The development of the 3C technique and its derivatives has provided a toolbox that enables the systematic spatial interrogation of multiple loci or even the entire genome(Lieberman-Aiden et al. 2009; Dekker et al. 2013). Based on genome-wide high-throughput chromosome conformation capture (Hi-C) technology, a 2D heat map can be constructed to indicate the frequency of interaction between two points in the genome. Depending on their positions, SVs often change the Hi-C profile of a locus, leaving specific signatures that can be analyzed further(Burton et al. 2013; Bianco et al. 2018). For example, deletions can result in novel interactions between two regions that were previously separated, whereas inversions result in a characteristic ‘bow tie’ configuration when mapped onto a reference genome. Furthermore, owing to its proximity-ligation nature, the Hi-C is also suitable for the identification of SVs without a priori knowledge(Rickman et al. 2012; Burton et al. 2013; Rao et al. 2014; Harewood et al. 2017).

SCRaMbLEing two synthetic chromosomes generated random and complex SVs, which reshaped chromosomes 3D architecture. Duan.et al reported that the genome 3D structure affected genes expressions(Duan et al. 2010) It is known that telomeres and centromeres could inhibit the expression of genes nearby (Chen and Zhang 2016), while the autonomously replicating sequence (ARS) could increase the expression of genes nearby(Flagfeldt et al. 2009). In the study, the pericentric inversions and trans- chromosomes translocation changed the map position of centromeres in chromosome X and caused the exchange of the right telomeres of chromosome V with chromosome X, respectively. Thus, larger number of genes position and the transcription direction, which affected the transcription levels of genes in the regions nearby the breakpoint of SVs. It is demonstrated that over-expression of YJR086W(STE18), YJR123W(RPS5) and YJR135C(MCM22) improved astaxanthin yield. Choudhury.et al show that phosphorylation of the yeast Gg subunit (Ste18) can serve as intrinsic regulators of G protein signaling and differentially activates mitogen-activated protein kinases (MAPKs) pathway (Choudhury et al. 2018), which potentially mediates the activation of some transcription factors and metabolic network related to the astaxanthin biosynthesis. Ghosh.et al reported Rps5 is an essential gene coding a protein component of the small (40S) ribosomal subunit. Over-expression of Rps5 potentially impact the translation initiation of yeast S. cerevisiae (Saliba et al. 2014), which may increase the translation of some genes being beneficial to the astaxanthin biosynthesis. Poddar.et al demonstrated MCM22 involved in chromosome segregation, mutations of which caused a decrease in the stability of the minichromosome . Over-expression of MCM22 potentially maintained the stability of chromosome during segregation and enhanced the cell viability during fermentation(Poddar et al. 1999). The exact mechanism of over- expression of STE18, RPS5 and MCM22 contributing to astaxanthin biosynthesis is unknown, however, upregulation of these genes may be beneficial for tuning metabolic network, the translation initiation process and stability of chromosome during segregation, which may explain why this genes affects astaxanthin biosynthesis.

In this manuscript, we have developed both a random method and a rational method to generate complex SVs. The random method uses SCRaMbLE in yeast with two synthetic chromosomes to generate yeast libraries containing complex SVs for screening desired phenotypes. The rational method enables a selection marker to switch from transcriptionally “off” to transcriptionally “on”. The rational method provides a convenient way to determine the effects of translocation and inversion targets on strain phenotypes. In addition, inversion and translocation strains generated from wild-type strains can be used as materials to study open genetic questions, such as whether inversion and translocation strains become reproductively isolated from control strains. By this method, we validated a larger fragment translocation (right arm of synV (YJR130C) and right arm of synX (YER164W)) and an inversion (YJL052C-A- YJR071C) in wild-type strains and found that both the translocation and inversion increased the yield of astaxanthin. This strategy is potentially a powerful tool for inducing the recombination of any two positions on chromosomes. How nature evolves with great biodiversity and nurtures organisms containing various numbers of chromosomes is not known. Our work may spark interest and provide a new handle for researchers to study this fundamental biological problem.

## Methods

### Strains and media

All yeast strains used in this study are described in Table 1. The strain SYNVX carries a synV, and a synX is derived from BY4741 (*MATa leu2Δ0 met15Δ0 ura3Δ0 his3Δ1*). BY4741 was used as the initial strain to reconstruct the translocations and inversions caused by SCRaMbLE of the strains and verify targets of astaxanthin yield improvement.

Yeast strains were cultured in YPD medium (10 g l^−1^ yeast extract, 20 g l^−1^ peptone, and 20 g l^−1^ glucose). SGal-Ura (synthetic media lacking uracil with 20 g l^−1^ galactose) with 1 μM β-estradiol was used to induce SCRaMbLE. SC medium containing 1 g L−1 5-FOA was used to screen strains without the URA3 marker. All yeast solid media contained 20 g L−1 agar. β-Estradiol and 5-FOA were purchased from Sigma-Aldrich. *E. coli* DH5α purchased from BEIJING Biomed Co., Ltd. was used for plasmid construction and replication. *E. coli* were cultivated at 37 °C in Luria–Bertani (LB) complete medium. Kanamycin (50 µg/mL) or ampicillin (100 µg/mL) was added to the medium for selection.

### Construction of plasmids and strains

YJJ001 (astaxanthin-producing control strain) was constructed by homologous recombination in SYNVX, directed by 500-bp CAN1 and 500-bp delta site genomic sequences flanking crtE-crtI-crtYB-crtZ-crtW-LEU2. YJJ002 was constructed by transforming YJJ001 with pCRE4 (Supplementary Fig. 1, Supplementary Data 2), followed by selection on SC-URA agar. Overexpression plasmids were constructed by Gibson assemble. The gene knockout cassette (left homologous arm-URA3-right homologous arm) was assembled by overlap extension PCR (OE-PCR).Transformations were performed using the standard lithium acetate procedure(Gietz et al. 1995).

### SCRaMbLE

YJJ002 containing the inducible Cre plasmid pGal1-Cre-EBD-tCYC1 was grown overnight in 5 mL SC-Ura media (30 °C, 250 r.p.m. shaking). Then, the cells were harvested and washed three times with sterile water to wash out glucose, and the culture was diluted to an OD600 of 0.6-1.0 in 3 mL SGal-Ura medium. Then, 1 μmol L^−1^ β- estradiol was added to the cultures to induce SCRaMbLE for 6 h (30 °C, 250 r.p.m. shaking). Cells were washed twice by centrifugation, resuspended in sterile water, diluted 5,000-fold, and plated on SC-Ura. The plates were incubated at 30 °C for 60 h.

### HPLC analysis of astaxanthin production

SCRaMbLE strains with darker colors were selected for fermentation in shake flasks. Three independent colonies of each strain were inoculated into 5 ml of YPD medium and grown at 30 °C until the OD 600≈ 8.0 (approximately 24 h). Then, the seed culture was transferred into 50 mL fresh YPD with 40 g l^−1^ glucose medium at an initial OD 600 of 0.1 and grown until ready to harvest.

Astaxanthin was extracted from HCl–heat treated cells with acetone according to Zhou et al. (Zhou et al. 2015) and Wang et al. (Wang et al. 2017) . Carotenoids were extracted as described below. Cells from 2 mL culture were collected and washed with distilled water. Then, the cells were resuspended in 1 mL of 3 M HCl, boiled for 5 min, and subsequently cooled in an ice bath for 5 min. After that, the cell debris was washed twice with distilled water and resuspended in 0.5 mL of acetone containing 1% (wv^−1^) butylated hydroxytoluene. Then, the mixture was vortexed until colorless (approximately 20 min) and incubated at 30 °C for 10 min. This was followed by centrifugation at 12,000 rpm for 5 min. The acetone phase containing the extracted astaxanthin was filtered through a 0.22-μm membrane for HPLC analysis.

### PCRTags analysis

All the synV and synX PCRTags involved in this study are listed in Supplementary Table 2. Amplification of PCRTags was performed using 7.5 μL 2 × Rapid Taq Master Mix (Vazyme Biotech Co., Ltd), 0.4 μL each of forward and reverse primers, 1 μL genomic DNA, and 5.7 μL ddH2O. The reaction procedure was as follows: 95 °C/1 min, 35 cycles of 95 °C/20 s, 53 °C/20 s, 72 °C/15 s and a final extension of 72 °C/5 min. Detection of PCRTags was performed by gel electrophoresis.

### Whole-genome sequencing (WGS) and analysis

WGS was performed at BGI (Beijing Genomic Institute in Shenzhen, China), and cells were harvested at exponential phase. Libraries were prepared and analyzed using an Illumina HiSeq X-Ten system. The sequencing data were filtered with SOAPnuke (v1.5.2)(Li et al. 2008). and clean reads were stored in FASTQ format for downstream analysis. The read comparison was performed using BWA software with the reference sequence. SVs including insertions, deletions, inversions, intrachromosomal translocations, and interchromosomal translocations were detected using Break Dancer software.

### Transcriptional analysis

Yeast cells were harvested from YPD medium at 24 h (exponential phase). Total RNA was extracted using the TRIzol® method following the NEB Next Ultra™ RNA protocol. The concentration of the extracted RNA samples was determined using a NanoDrop system (NanoDrop, Madison, USA), and the integrity of the RNA was examined based on the RNA integrity number (RIN) determined using an Agilent 2100 Bioanalyzer (Agilent, Santa Clara, USA). RNA sequencing was carried out by the BGISeq500 platform (BGI-Shenzhen, China). The sequencing data were filtered with SOAPnuke (v1.5.2).(Li et al. 2008)

Clean reads were obtained and stored in FASTQ format. The clean reads were mapped to the reference genome using HISAT2 (v2.0.4)(Kim et al. 2015). Bowtie2 (v2.2.5)(Langmead and Salzberg 2012) was applied to align the clean reads to the reference coding gene set, and the expression level of each gene was calculated by RSEM (v1.2.12)(Li and Dewey 2011). The heatmap of the gene expression in different samples was drawn by GraphPad Prism (v8.0.1). Essentially, differential expression analysis was performed using DESeq2 (v1.4.5)(Love et al. 2014) with a Q value ≤ 0.05. To gain insight into the change in phenotype, GO (http://www.geneontology.org/) and KEGG (https://www.kegg.jp/) enrichment analyses of annotated differentially expressed genes were performed by Phyper (https://en.wikipedia.org/wiki/Hypergeometric_distribution) based on the hypergeometric test. Significant terms and pathways were identified as those with Bonferroni-corrected *P*-values below a rigorous threshold (*P*-value ≤ 0.05). Triplicate samples were used for transcriptional analysis. The Saccharomyces Genome Database (SGD)(Cherry et al. 2012) was used to obtain gene information.

### Hi-C library generation and sequencing

Hi-C library generation and sequencing were carried out by Frasergen (Wuhan, China), and cells were harvested at exponential phase. Cells were resuspended in 1×PBS to an OD600 of 1.0. Cells were cross-linked in a 3% final concentration of fresh formaldehyde and quenched with glycine (0.15 M final concentration) for 5 min. The cells were resuspended in 1 mL of 1× NEBuffer 2.1 (NEB) and homogenized by grinding to a fine powder in liquid nitrogen. Then, the homogenized yeast material was washed with 25 mL of 1× NEBuffer 2.1 and suspended in 2.5 mL of 1× NEBuffer 2.1. Cells were split into aliquots (V =456 µL) and solubilized in 0.1% SDS for 10 min at 65 °C. Cross-linked DNA was digested with 200 U MboI (NEB) per tube at 37 °C overnight. Restriction fragment ends were labeled with biotinylated cytosine nucleotides by biotin-14-dCTP (TriLINK). Blunt-end ligation was carried out at 16 °C overnight in the presence of 100 Weiss units of T4 DNA ligase (Thermo, 10.0 mL final volume per tube).

DNA purification was achieved through overnight incubation at 65 °C with 200 µg/mL proteinase K (Thermo). Purified DNA was sheared to a length of ∼400 bp. Point ligation junctions were pulled down by Dynabeads® MyOne™ Streptavidin C1 (Thermo Fisher). The Hi-C library for Illumina sequencing was prepared using the NEBNext® Ultra™ II DNA library Prep Kit for Illumina (NEB) according to the manufacturer’s instructions. Fragments of between 400 and 600 bp were paired-end sequenced on an Illumina HiSeq X10 platform (San Diego, CA, United States) in 150PEmode. Two replicates were generated for each group of materials.

### Construction of contact map

The contact maps were generated using the ICE software package (version 1f8815d0cc9e)(Imakaev et al. 2012), and the Hi-C data of YJJ001, YJJ168 and YJJ432 cells were iteratively mapped to their own genomes. Dangling ends and other unusable data were filtered out, and the valid pairs were binned into 10 kb nonoverlapping genomic intervals to generate contact maps. The contact maps were normalized using an iterative normalization method to eliminate systematic biases.

### Cre/loxP induced chromosome translocation and inversion

Translocations and inversions were detected in YJJ168 and YJJ432 and verified in the wild-type strain BY4741 by introducing two Cre/loxP sites to the corresponding position. LoxP sites were integrated by homologous recombination. Homologous arms upstream and downstream were amplified from the genome of BY4741. Two loxP sites (loxP site 1 and loxP site 2) were integrated along with URA3+ and HIS3+/Hyg- as selection markers. The URA3 promoter was inserted upstream of loxP site 1, and open reading frames of URA3 were inserted downstream of the loxP site, allowing the expression of URA3. Hyg without a promoter was positioned adjacent to downstream loxP site 2 and the HIS3 promoter, and open reading frames of HIS3 were positioned upstream of loxP site 2. Adding 1 μmol L–1 β-estradiol to the cultures to switch on Cre-mediated recombination between the two loxP sites results in translocation and inversion, which subsequently turn on the expression of the Hyg gene and simultaneously shut down URA3 expression. Translocation and inversion strains could be selected on SC-LEU-5-FOA plates and SC-LEU+Hyg.

### Data availability

The data that support the findings of this study are available from the corresponding author on request. Transcriptomes data and Whole-genome sequencing data are available at Sequence Read Archive (SRA) under accession code SUB8866811.

## Acknowledgements

This work was funded by Ministry of science and technology the National Key Research and Development Program of China (2021YFC2100800), and the National Natural Science Foundation of China (31800719, 31861143017 and 21621004).

## Author contributions

B.J., and J.J. contributed equally to this work. B.J., J.J., and Y.J.Y. designed the experiments. B.J., J.J., and M.Z.H performed the experiments. B.J., J.J., and Y.J.Y. wrote the manuscript and all authors edited the manuscript. This project was supervised by Y.J.Y.

