## Supplemental information for "Directed yeast genome evolution by controlled introduction of trans-chromosomic structural variations"

**Supplementary Table S1.** Plasmids used in this study

| Plasmid | Description | Sources |
|---|---|---|
| pAsta | pRS416, pTEF1-crtE-tPDX1-pTDH3-crtI-tMPE1-pFBA1-crtYB-tTDH2-ADH1t-Aa_crtZ-FBA1p-TDH3p-BDC263_crtW-TDH2t | This study |
| pCRE4 | pGAL1-Cre-EBD-CYC1t | Jia. et.al |
| pCRE7 | pGAL1-Cre-EBD-CYC1t-G418 | This study |
| pJJ522 | pRS416, pTDH3-YJL053W-tADH1 | This study |
| pJJ523 | pRS416, pTDH3-YJL052C-A-tADH1 | This study |
| pJJ525 | pRS416, pTDH3-YJL049W-tADH1 | This study |
| pJJ526 | pRS416, pTDH3-YJL046W-tADH1 | This study |
| pJJ527 | pRS416, pTDH3-YJL045W -tADH1 | This study |
| pJJ528 | pRS416, pTDH3-YJR067C-tADH1 | This study |
| pJJ529 | pRS416, pTDH3-YJR068W-tADH1 | This study |
| pJJ530 | pRS416, pTDH3-YJR070C-tADH1 | This study |
| pJJ531 | pRS416, pTDH3-YJR071W-tADH1 | This study |
| pJJ532 | pRS416, pTDH3-YJR072C-tADH1 | This study |
| pJJ533 | pRS416, pTDH3-YJR076C-tADH1 | This study |
| pJJ534 | pRS416, pTDH3-YJR079W-tADH1 | This study |
| pJJ535 | pRS416, pTDH3-YJR080C-tADH1 | This study |
| pJJ536 | pRS416, pTDH3-YJR082C-tADH1 | This study |
| pJJ537 | pRS416, pTDH3-YJR085C-tADH1 | This study |
| pJJ538 | pRS416, pTDH3-YJR086W-tADH1 | This study |
| pJJ539 | pRS416, pTDH3-YJR087W-tADH1 | This study |
| pJJ540 | pRS416, pTDH3-YJR088C-tADH1 | This study |
| pJJ541 | pRS416, pTDH3-YER156C-tADH1 | This study |
| pJJ543 | pRS416, pTDH3-YER163C-tADH1 | This study |
| pJJ544 | pRS416, pTDH3-YER168C-tADH1 | This study |
| pJJ545 | pRS416, pTDH3-YER170W-tADH1 | This study |
| pJJ546 | pRS416, pTDH3-YJR122W-tADH1 | This study |
| pJJ547 | pRS416, pTDH3-YJR123W-tADH1 | This study |
| pJJ549 | pRS416, pTDH3-YJR125C-tADH1 | This study |
| pJJ551 | pRS416, pTDH3-YJR133W-tADH1 | This study |
| pJJ552 | pRS416, pTDH3-YJR135C-tADH1 | This study |
| pJJ553 | pRS416, pTDH3-YJR135W-A-tADH1 | This study |
| pJJ554 | pRS416, pTDH3-YJR136C-tADH1 | This study |

**Supplementary Table S2.** Primers used in this study

| Primer | Sequence (5'- 3') |
|---|---|
| **For replacement of *YJR116W*** | |
| YJR116W_LF | tacgacgcggatctgcccgtcatcgcatac |
| YJR116W_LR | AAAAAAAATGATGAATTGAAttcagcagtgggtcacggat |
| URA3 _F | atccgtgacccactgctgaaTTCAATTCATCATTTTTTTT |
| URA3 _R | ctgtgtaggtgacaacattaTTAGTTTTGCTGGCCGCATC |
| YJR116W_RF | GATGCGGCCAGCAAAACTAAtaatgttgtcacctacacag |
| YJR116W_RR | ctagcttcgaggtaggagcaggcttagcca |
| **PCR verification of replacement of *YJR116W*** | |
| YJR116W_RVF1 | agatacgttccaagctgtct |
| YJR116W_RVR1 | TGCCCATTCTGCTATTCTGT |
| YJR116W_RVF2 | ACAGAATAGCAGAATGGGCA |
| YJR116W_RVR2 | ggatggtcgagaatcgtctt |
| **PCR verification of deletion of *YJR116W*** | |
| YJR116W_DVF1 | ataccggtgatcaagtctca |
| YJR116W_DVR1 | ttaggatggtcgagaatcgt |
| **PCR verification of junction of translocation in YJJ432** | |
| SynV_VF1 | cacgagattgtctccaaaca |
| SynV_VR1 | cagaaacctctttccttgga |
| SynX_VF2 | ttgacaattgtgaatgcctt |
| SynX_VR2 | aggataccaagcggaattat |
| **PCR verification of junction of inversion in YJJ168** | |
| YJR071W _VF1 | agctaagtaaaagcagcttggag |
| YJR071W _VR1 | gttgattctgttaatggccag |
| YJL052C _VF2 | tggtaggtagcgatcttgac |
| YJL052C _VR2 | agaactctgctccagaccta |
| **PCR verification of junction of translocation in YJJ474** | |
| SynV_VF1 | cacgagattgtctccaaaca |
| SynV_VR1 | cagaaacctctttccttgga |
| SynX_VF2 | ttgacaattgtgaatgcctt |
| SynX_VR2 | aggataccaagcggaattat |
| **PCR verification of junction of inversion in YJJ473** | |
| SynV_VF1 | agctaagtaaaagcagcttggag |

| | |
|---|---|
| SynV_VR1 | gttgattctgttaatggccag |
| SynX_VF2 | tggtaggtagcgatcttgac |
| SynX_VR2 | agaactctgctccagaccta |

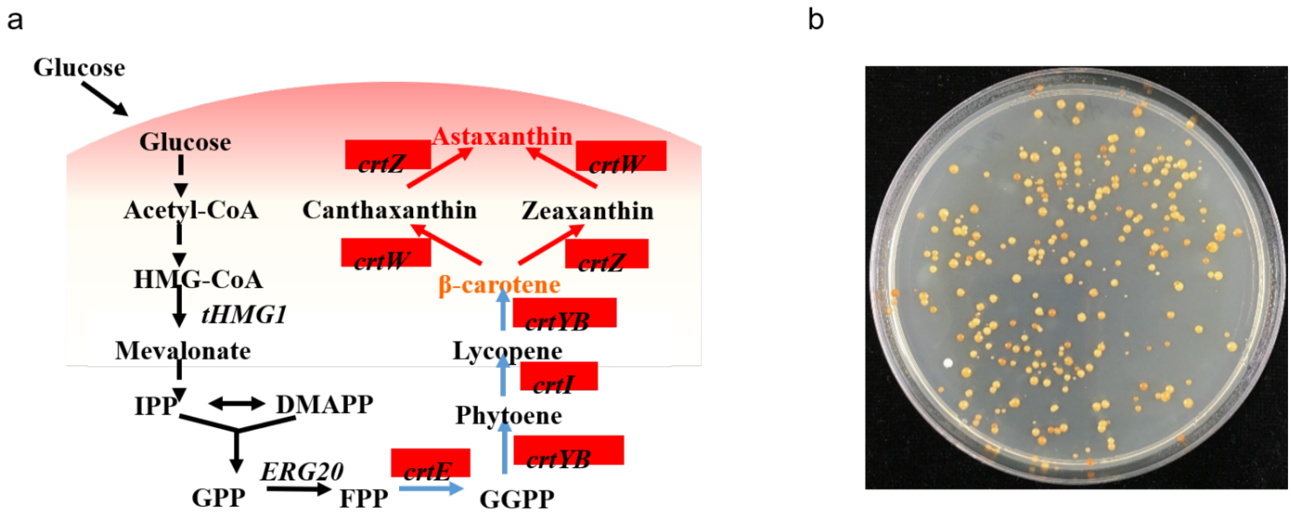


**Supplementary Figure S1.** a. Biosynthesis pathway of astaxanthin in yeast. b. SCRaMbLEd yeast pool. The SCRaMbLE library was plated on SC-Ura glucose agar and the strains incubated at 30°C for 60 h.

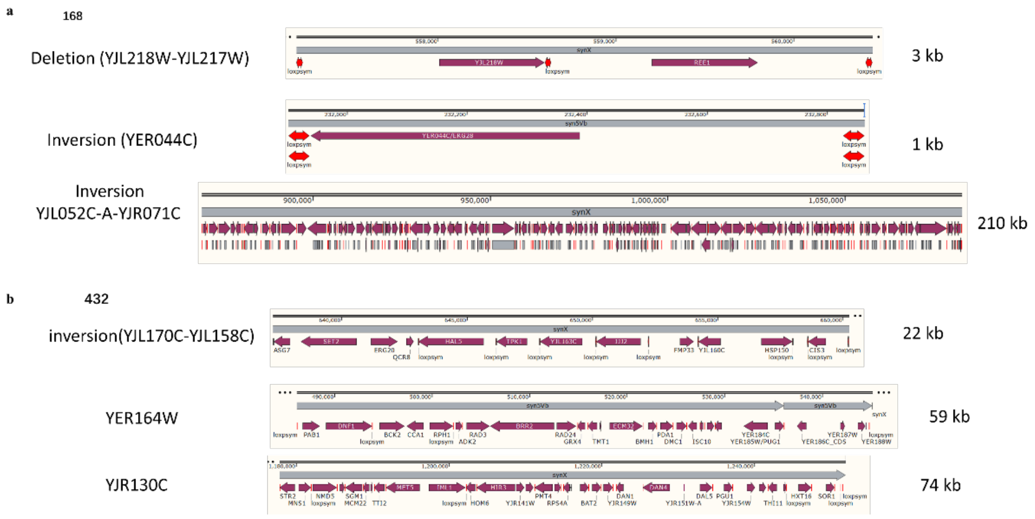


**Supplementary Figure S2.** Sequence analysis of YJJ168 and YJJ432. a 3 kb deletion (YJL218W-YJL217W), a 1 kb inversion (YER044C) and a 210 kb inversion (YJL052C-A-YJR071C) were observed in the YJJ168. a 22 kb inversion (YJL170C-YJL158C) and a translocation (the 59 kb YER164W-a 74 kbYJR130C) were observed in the YJJ432.

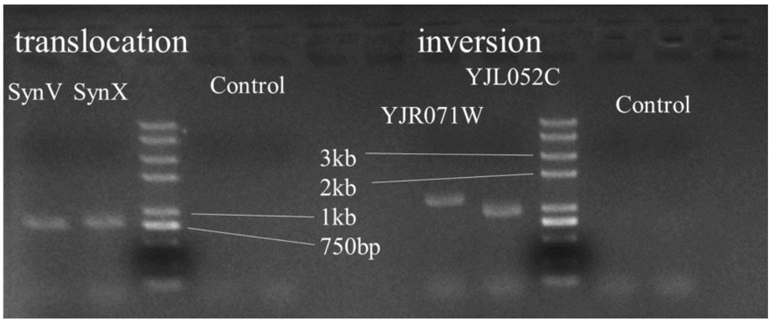


**Supplementary Figure S3.** PCR verification of inversion strain yJJ168 and translocation strain yJJ432. To be notably, the primers could amplify the band from control, which should be 768 bp (for synV of translocation in yJJ432), 788bp (for synX of translocation in yJJ432), 1190 bp (for YJR071W of inversion in yJJ168 and yJJ432) and 951 bp (for YJL052C of inversion in yJJ168 and yJJ432).

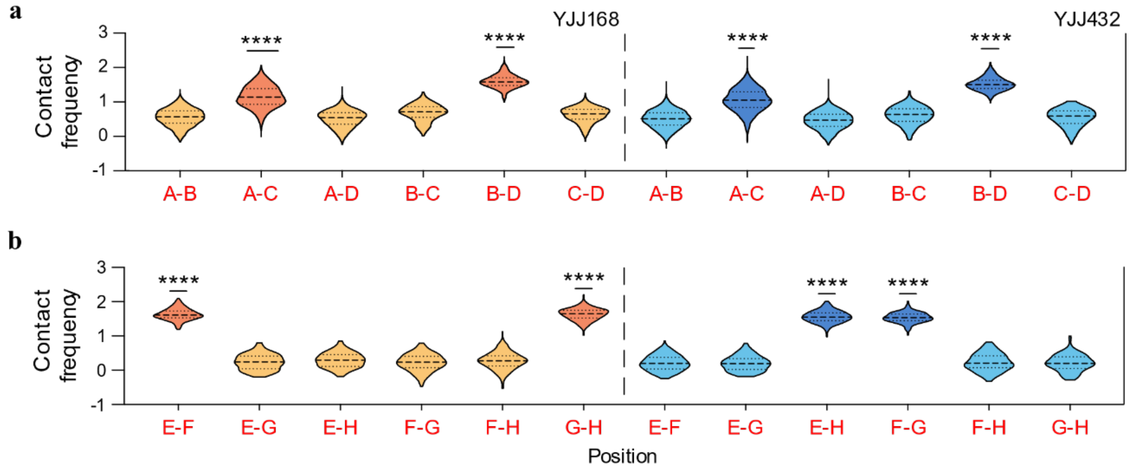


**Supplementary Figure S4.** Violin plot of the contact frequencies between recombination sites. ****P < 0.0001.

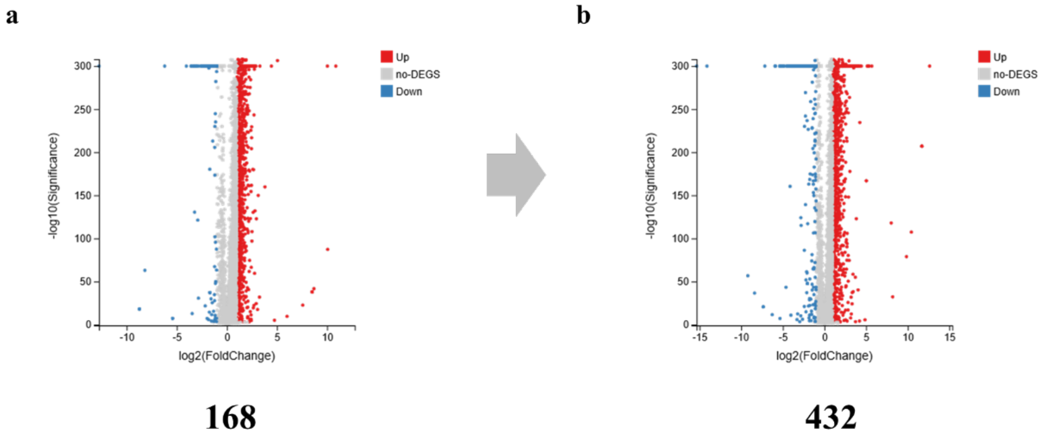


**Supplementary Figure S5.** The volcano plot of transcriptome of yJJ168 (a) and yJJ432 (b). No-differentially expressed genes are shown in gray, up expressed genes are shown in red and down genes are shown in blue.

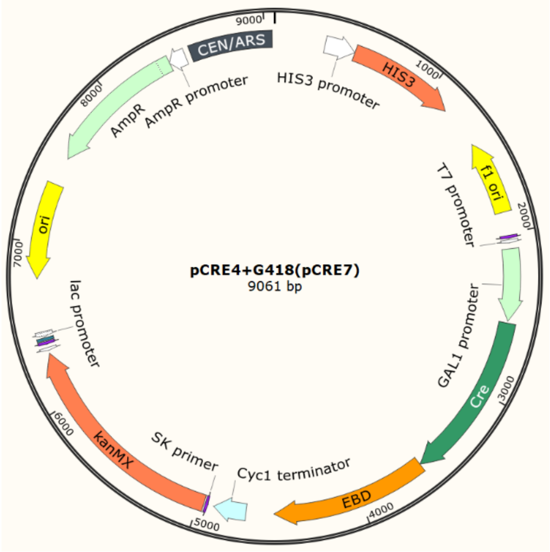


**Supplementary Figure S6.** Plasmids map of CRE switches pCRE7: pGAL1-Cre-EBD-tCYC1-G418 used in this study.
